# Tipping points emerge from weak mutualism in metacommunities

**DOI:** 10.1101/2023.02.26.530120

**Authors:** Jonas Denk, Oskar Hallatschek

## Abstract

The coexistence of obligate mutualists is often precariously close to tipping points where small environmental changes can drive catastrophic shifts in species composition. For example, microbial ecosystems can collapse by the decline of a strain that provides an essential resource on which other strains cross-feed. Here, we show that tipping points, ecosystem collapse, bistability and hysteresis arise even with very weak (non-obligate) mutualism provided the population is spatially structured. Based on numeric solutions of a metacommunity model and mean-field analyses, we demonstrate that weak mutualism lowers the minimal dispersal rate necessary to avoid stochastic extinction, while species need to overcome a mean threshold density to survive in this low dispersal rate regime. Moreover, we show that, starting with a randomly interacting species pool, metapopulation structure tends to select for an ecosystem with mutualistic interactions. Bistable metacommunities could, therefore, be a natural outcome of evolutionary dynamics in structured ecosystems.

## Introduction

Mutualistic interactions between species are ubiquitous in nature and can be critical for the stability of natural ecosystems as exemplified by cross-feeding microbes in the gut (1–3), cooperative growth (4), and sexual mating (5). When mutualism is obligate, i.e. species survival relies on the presence of each other, the well-mixed population dynamics can exhibit a critical threshold population size that a species must overcome to avoid extinction, often referred to as strong Allee effect (6, 7) [see Fig. 1 (a)]^1^. Models that incorporate a strong Allee effect are of great interest in ecology and are invoked frequently to explain tipping points and catastrophic shifts between survival and extinction in ecosystems (8–10). As an instructive example, when microbes depend on each other in order to access vital resources, their populations dynamics can exhibit a tipping point at which the community undergoes catastrophic shifts upon the variation of experimental parameters such as nutrient levels (4, 11–13)

**Fig. 1.**
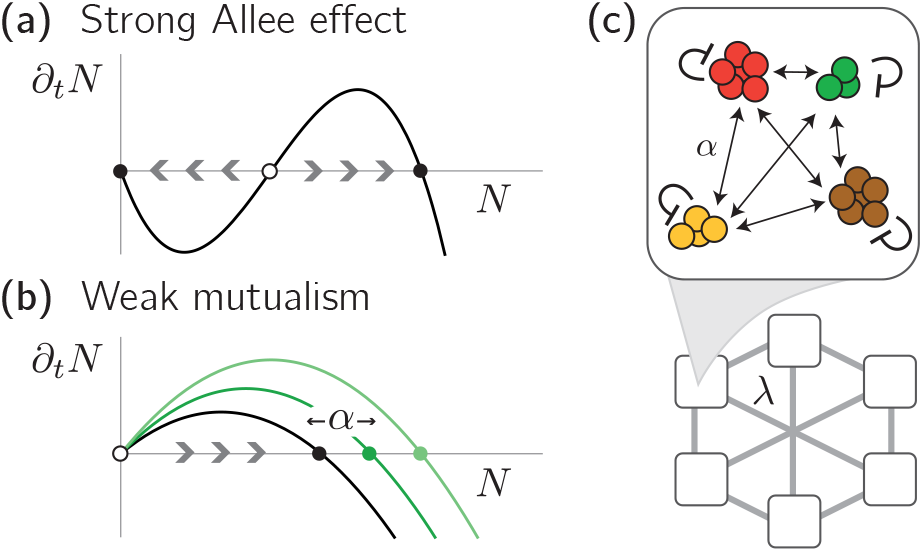
Obligate and weak mutualism. **(a)** Obligate mutualism can lead to a strong Allee effect, where the dynamics *∂_t_N* of a well-mixed population of size *N* leads to either extinction (*N* = 0)or a finite population size (*N* > 0) depending on the initial population size. Full and open circles denote stable and unstable fixed points, respectively, where the center unstable fixed point is often referred to as Allee threshold. Arrows denote the flow of the dynamics. **(b)** In contrast, weak mutualism merely increases the effective carrying capacity (stable fixed points) of a species, which is achieved by increasing *a* in our model. **(c)** Illustration of our mathematical approach. Different species (different colors) interact through weak mutualism with a strength *a* and all individuals disperse between all patches at a dispersal rate *λ* (global dispersal).

When populations are coupled through dispersal, i.e. in a metapopulation, bistability of the respective well-mixed population dynamics can lead to spatiotemporal behavior that is qualitatively different than in a metapopulations undergoing regular logistic growth, including pushed instead of pulled waves in range expansions (14–16), localized wave fronts (17), and pronounced patchiness (18).

As opposed to obligate mutualism, we will refer to ecosystems with *weak* mutualism as ecosystems whose well-mixed dynamics show neither tipping points nor bistability, but follow simple logistic growth qualitatively similar to a population in the absence of inter-species interactions [see Fig. 1(b)]. Accordingly, it is less clear how, if at all, a metacommunity with weak mutualism shows different spatiotemporal dynamics than a metacommunity without any interspecies interactions.

Here, we show that even the weakest form of mutualism in a metacommunity can lead to a tipping point accompanied by bistability and catastrophic shifts between coexistence and extinction when demographic fluctuations are taken into account. Our combined analytical and numerical methods reveal a regime of intermediate dispersal rates, where species can avoid extinction when the mean population size overcomes a threshold size. Informed by our intuition regarding purely mutualistic interactions, we show that close to the onset of finite population sizes, metacommunities with random interactions undergo selection for mutualistic interactions. We further apply our analyses to metacommunities in which mutualism acts on the dispersal rate rather than inter-species interactions and find a similar emergent tipping point. Our results give insights into the role of demographic stochasticity and dispersal in metacommunities and highlight the emergence of tipping points and catastrophic shifts even when absent under well-mixed conditions.

## Results

### Mathematical approach to metaeommunities with weak mutualism

In the following, we consider *S* species that live in a metacommunity of *P* coupled communities (patches), where *P* is assumed to be large. The dynamics of the population size *N_x,i_* of species *i* ∈ {1,...*S*} on patch *x* ∈ {1,...,*P*} is modeled by the following set of generalized Lotka-Volterra equations [see Fig. 1(c) for a graphical representation]:

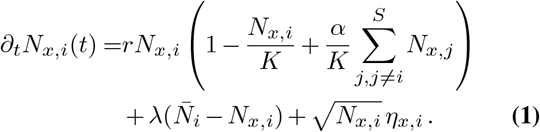

The first term in Eq. (1) describes growth of a species’ population at a growth rate *r* > 0, which saturates at a carrying capacity *K* due to self-limiting interactions. The interaction parameter *α* > 0 denotes the strength of mutualistic interactions between species. Assuming a constant interaction strength *α* between all species allows an analytic meanfield description; the results of this analysis will yield important intuitions when we later allow variations in the species’ inter-species interactions. The second term in Eq. (1) takes into account dispersal, where we assumed, as a simple spatial approach, that all patches are connected through dispersal with a species-independent dispersal rate *λ* [global dispersal, compare Fig. 1(c)], and 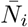 denotes the abundance of species *i* averaged over all *P* patches. The last term in Eq. (1) reflects demographic fluctuations due to random births and deaths of individuals within a population, where *η_x,i_* denotes uncorrelated noise with zero mean and variance *ω*^2^. The square-root dependence of demographic noise on the density ensures that the expected variance of fluctuations is proportional to the expected number of birth or death events during one generation and has been derived in various contexts from discrete descriptions of growing populations (19, 20). Under well-mixed conditions, the deterministic population dynamics of Eq. (1) for every species follows regular logistic growth with a stable steady state abundance *N\** at

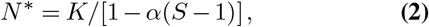

as long as mutualistic interactions are weaker than self-limiting interactions, i.e. *α* < (*S* – 1)^-1^. Specifically, this means, that population growth does *not* display an explicit Allee effect nor bistability [compare Fig. 1(a) and (b)]. In the following, we will solve Eq. (1) numerically and employ mean-field analyses to study the effect of demographic fluctuations and dispersal and show how these can, nevertheless, generate bistability and an abrupt shift in the population size.

### Weak mutualism generates a tipping point in a metacommunity

For a clearer presentation of our results, in the following we fix *r, K*, and *α* (with *α* ≪ (*S* – 1)^-1^), and vary the dispersal rate *λ* for different numbers of species *S*. First, we discuss our numerical solutions of Eq. (1) assuming small average abundances *N* = (*SP*)^-1^ ∑_*x i*_ *N_x,i_* as initial condition (for details on the numerical solution, see Appendix 1). In the absence of mutualistic interactions, i.e. *S* = 1, the growth and dispersal dynamics represented by Eq. (1) have been extensively studied for short-range dispersal in the context of directed percolation (21, 22) and for global dispersal in metapopulations (23–26). From these earlier studies we expect that for *S* = 1, increasing the dispersal rate leads to a continuous transition from a phase of zero population size (absorbing phase) to a phase of finite population sizes (active phase). Indeed, when numerically solving the dynamics Eq. (1) with global dispersal and for only one species (*S* = 1), we find that for zero and small dispersal rates *λ* all species eventually go extinct, whereas for *λ* above a critical threshold value *λ_c_*, the average population size after the final time step of our numerical solution is finite and increases continuously with *λ* [see triangles in Fig. 2(a)]. Interestingly, when increasing the number of species at constant mutualistic interaction strength *α* > 0, the average abundance as a function of the dispersal rate *λ* undergoes a sudden jump at *λ_c_* from zero to finite values. Discontinuous transitions are a telltale sign of subcritical bifurcations and bistability close to the transition (27). To investigate the possibility of multiple stable solutions, we repeat our numerical solutions for larger initial average abundances, and, indeed, find bistability close to the threshold dispersal rate *λ_c_* for larger *S* [see circles in Fig. 2(a)].

**Fig. 2.**
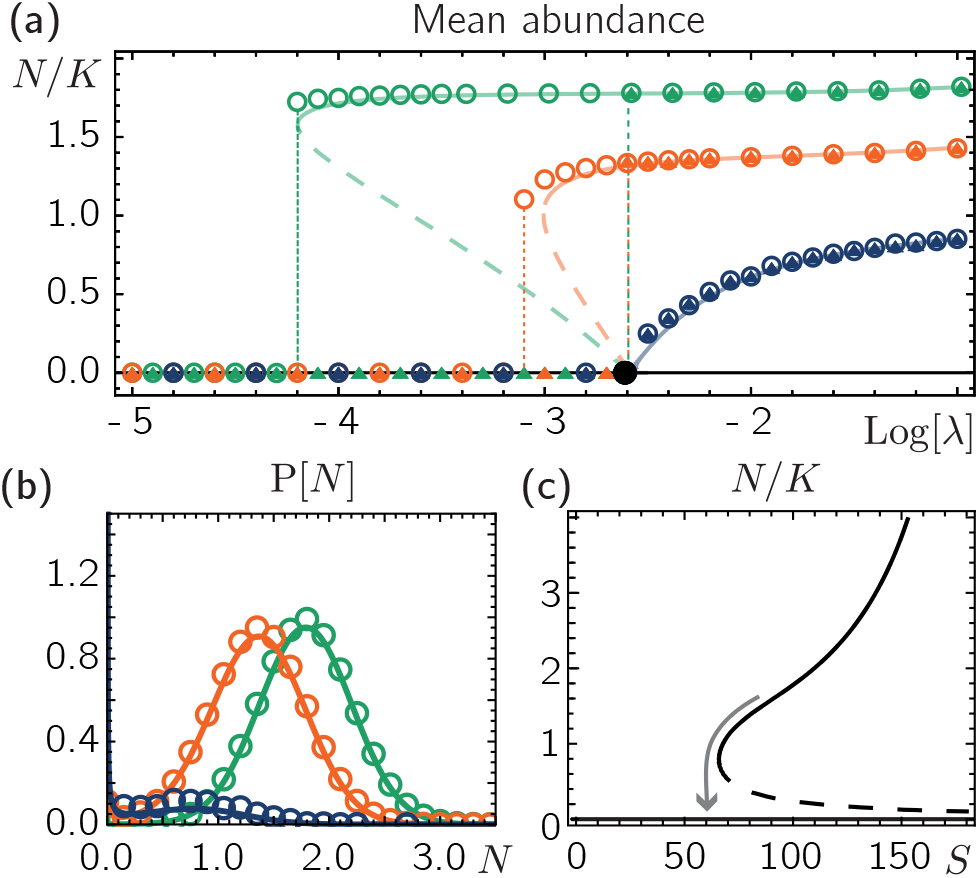
Weak mutualism generates a tipping point. **(a)** Starting at small and large initial population sizes (triangles and circles, respectively), the mean population size in our numerical solutions can reach zero or finite values. These numerical results are in very good agreement with our mean-field solution (lines). Solid and dashed lines denote stable and unstable manifolds, respectively. Colors denote different numbers of species, *S* = 100 (green), *S* = 75 (orange), and *S* =1 (blue). The threshold dispersal rate *λ_c_* is shown as a full black circle. **(b)** Numerical and analytic solutions for the abundance distribution *P*[*N*] (circles and lines, respectively) for dispersal rates just above *λc*. Colors represent different *S* as in (a). **(c)** Small changes in the species number, i.e. through perturbations, can lead to a collapse of the metacommunity (as indicated by the arrow), *λ* = 0.001. Parameters: *r* = 0.3, *K* = 10, *α* = 0.005, *P* = 500.

To get deeper insights into the exact form of the bifurcation, we employ a mean-field approximation of Eq. (1), that has been recently presented to approximate the stationary abundance distribution for metacommunities with competitive interactions (23). In short, by expressing the interaction term through the species-averaged abundance on a patch defined as 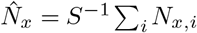 and treating the mean-fields 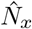 and 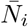 as deterministic mean-field parameters, we can map the dynamics Eq. (1) to the solvable problem of a Brownian particle in a fixed potential. Demanding that 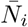 and 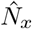 are both equal in equilibrium (in the limit of an infinite number of species and patches) we can derive an analytic expression for the abundance distribution as a function of the mean species abundance 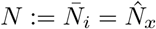 and the control parameters *r, K, α, S*, and *λ*, which can be solved self-consistently [for a detailed derivation see (23) and Appendix 2]. In agreement with our numerical solution, our analytic mean-field approximation predicts bistability between the dispersal rate *λ_c_* [for an analytic expression see (23) and Appendix 2] and a lower dispersal rate *λ_t_*, which marks a point where a small decrease of the dispersal rate causes an abrupt shift from finite population size to extinction, often referred to as tipping point (10) [see dashed vertical lines in Fig. 2(a)]. In addition, our analytic solution reveals an unstable branch marking the threshold mean abundance as a function of the dispersal rate the metacommunity must exceed to reach a finite mean abundance and avoid extinction [see long dashed lines in Fig. 2(a)]. Looking at the abundance distribution 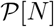 at a dispersal rate close above *λ_c_*, we observe that when increasing the number of species, its shape transitions from a scaling 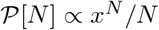 for small *N*, a common scaling in ecology often referred to as Fisher-log series (28, 29), to an approximate Gaussian distribution [see Fig. 2(b), for details see (23) and Appendix 2].

In summary, our numerical and analytical results strongly suggest that despite the lack of bistability in the deterministic and well-mixed population dynamics, the metacommunity displays a tipping point accompanied by a regime of bistability. The identified bifurcation predicts discontinuous transitions at *λ_c_* and the tipping point dispersal rate *λ_t_*, and suggests hysteresis when varying the dispersal rates across these two values. We find that the range of bistability increases the more species interact through mutualism [see Fig. 2(c)]. As a consequence, this suggests that perturbations that decrease the number of species, even if only by a few species, may shift the metacommunity into a regime where eventually all species go extinct [see arrow in Fig. 2(c)]. While we find that demographic fluctuations can lead to bistability, even when bistability is absent under deterministic well-mixed conditions, previous studies suggest that demographic fluctuations can also have the reverse effect on population transitions: a metapopulation that exhibits bistability under deterministic and well-mixed conditions (imposed through an explicit strong Allee effect) can show a smooth transition when dispersal and stochasticity are taken into account [see (30–32) and Appendix 3]. Comparing these previous results with our results for weak mutualism (Fig. 1) highlights the multifaceted role of stochasticity in spatially extended populations.

### Spatial structure selects for metacommunities with an excess of mutualistic interactions

The dramatic change from a smooth to a discontinuous transition close to the threshold dispersal rate *λ_c_* suggests several implications for more general species interactions. Since in contrast to species with competitive interactions, species with mutualistic interactions are able to reach large finite abundances already well below *λ_c_*, we hypothesize that in a metacommunity with random interactions, mutualism may play an important role in community assembly, at least close to *λ_c_*. To test this idea, we generalize the metacommunity dynamics Eq. (1) and assume random interactions *α_i,j_* between species *i* and *j*. The generalized metacommunity dynamics reads

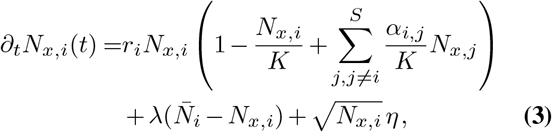

where we again assumed global dispersal between patches. For a clearer distinction between mutualistic and competitive interactions between species, we choose interactions to be symmetric, i.e. *α_j,i_ = α_i,j_*. Motivated by earlier work on well-mixed communities (33–38), we draw *α_i,j_*, from a Gaussian distribution with mean *α* and standard deviation 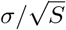. Hence, it is possible to choose *α* and *σ* in a way that interactions between some species *i* and *j* are mutualistic (*α_i,j_* > 0) while interactions are on average competitive (*α* < 0). A detailed analytical understanding of the spatially structured community assembly with random interactions is beyond the scope of our mean field analysis, but can be obtained using the replica method (39). In the following we use numerical solutions of Eq. (3) to show that, below the critical threshold dispersal rate *λ_c_*, communities survive that are enriched in mutualistic interactions, even though the interactions among the initial species pool are on average competitive.

First, we solve the dynamics Eq. (3) numerically for fixed *r, K, S, σ*, negative interaction mean *α* (i.e. an on average competitive interaction between species), and varying *λ*. Here, we will focus on relatively small interaction differences, i.e. small *σ*, where previous studies suggest that, under well-mixed conditions, a community approaches a unique equilibrium state (35–37). Similar to our results for purely mutualistic interactions [Fig. 2(a)], we find positive population sizes already for dispersal rates below *λ_c_* [see Fig. 3(a)]. Furthermore, we observe bistability, i.e. for dispersal rates below *λ_c_* the metacommunity approaches positive mean population sizes when the initial population size is sufficiently large while it goes extinct otherwise. In order to investigate the role of mutualistic interactions especially in this regime of dispersal rates around *λ_c_*, we calculate the mean interaction *I*: = 〈(*α_i,j_*/*K*)*N_j,x_*〉_*x,j*_ of all surviving species at the last time point of our numerical solution [see Fig. 3(b)]. In line with our intuition from purely mutualistic interactions, we find that for dispersal rates below *λ_c_*, all species that survive form on average mutualistic interactions with their fellow surviving species [the distribution of *I*, 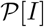, in Fig. 3(b) has only contributions from positive *I*]. When we increase the dispersal rate beyond *λ_c_*, more species survive and, in particular, also species with on average competitive interactions coexist. We were also interested how the communities of remaining survivors of our numerical solution compare to our mean-field model with species-independent interaction coefficient. To this end, we calculate the mean-field solution with the number of species *S* and the species-independent interaction coefficient *α* being equal to the number of surviving species and their average interaction coefficient, respectively, that we obtained from our numeric simulation of the metacommunity with random interactions. Interestingly, we observe that in the bistability regime, and when starting at large population sizes, the surviving community is very close to the tipping point of our mean-field prediction (see Appendix 4). This suggests that random metacommunities self-organize to a state very close to the tipping point where they may be very sensitive towards perturbations in their parameters e.g. their number of species.

**Fig. 3.**
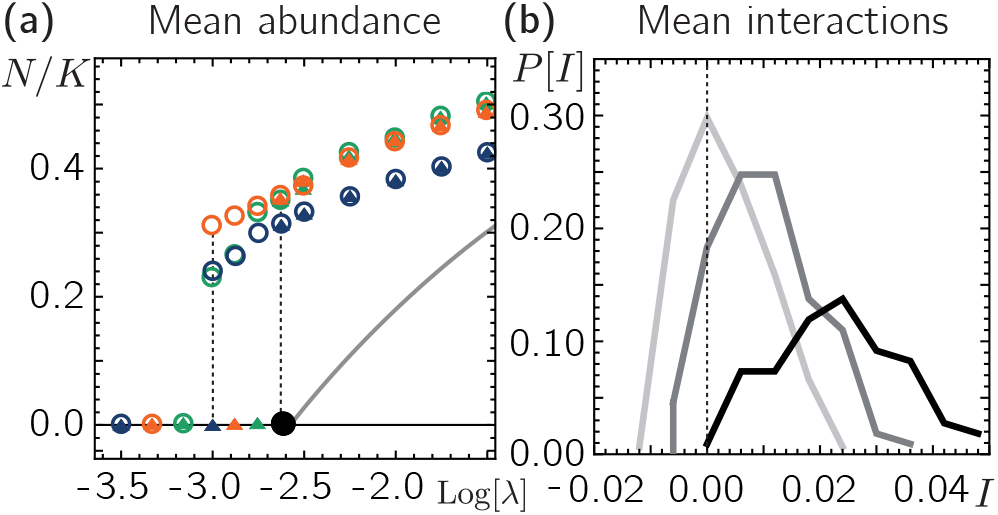
Tipping point for metacommunities with random interactions. **(a)** Numerical solutions of Eq. (3) for initially low (triangles) and high (circles) mean population sizes for three different sets of random interactions (denoted by three different colors) suggest bistability and hysteresis between the tipping point (left dashed line) and close to *λ_c_* (black circle, right dashed line). Gray solid line shows mean-field solution for identical, competitive interactions (*α_i,j_* = *α*) **(b)** Distributions of mean interactions with other species for all three sets of random interactions shown in (a) for *λ* = 10^-2.8^, *λ* = 10^-2.2^, *λ* = 10^-1.5^ from light to dark, respectively (*λ_c_* ≈ 10^-2.6^). Remaining Parameters: *σ* = 0.5,*r* = 0.3,*S* = 100,*α* = −0.01, *P* = 500.

### Tipping point through density-dependent dispersal

So far, we have incorporated mutualism through direct interactions between species, such that interactions effectively increase the species growth [see Eq. (1) and Eq. (3)]. In the following, we investigate positive interactions between species through their dispersal, i.e. interactions that increase the species’ dispersal rates.

There is indeed evidence in many species (40–44) that emigration rates from crowded areas tend to be elevated to avoid competition for resources. Since this effectively results in a dispersal rate that increases with the abundance of other populations, we refer to this scenario as mutualism through density-dependent dispersal in the following. When assuming that, as a first approximation, emigration from a patch increases linearly with the number of individuals of other species already present, the dispersal term in Eq. (1) can be written as

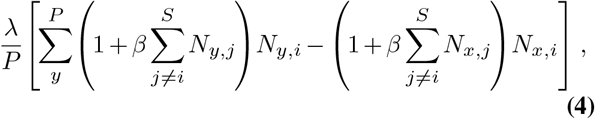

where we assumed a constant baseline dispersal rate *λ* between every patch and a linear increase of dispersal with population size with a factor *β*. Next, we solve Eq. (1) for fixed *r*, *K*, and *S* > 1 with *α* = 0, i.e. without direct mutualistic interactions, and the dispersal term Eq. (4) with *β* > 0 numerically and discuss the respective mean-field approximation (for details on the mean-field description, see Appendix 2). Similarly to direct mutualistic interactions between species, we find that when varying the baseline dispersal rate *λ*, the average abundance of the metacommunity undergoes a sub-critical bifurcation at *λ_c_* [see Fig. 4(a)]. As for direct mutualistic interactions, the regime of dispersal rates *λ* that shows bistability grows for increasing *S* [see Fig. 4(b)]. We thus conclude that mutualistic interactions through density-dependent dispersal result in a similar phenomenology, including catastrophic shifts, as mutualism that directly affects species growth.

**Fig. 4.**
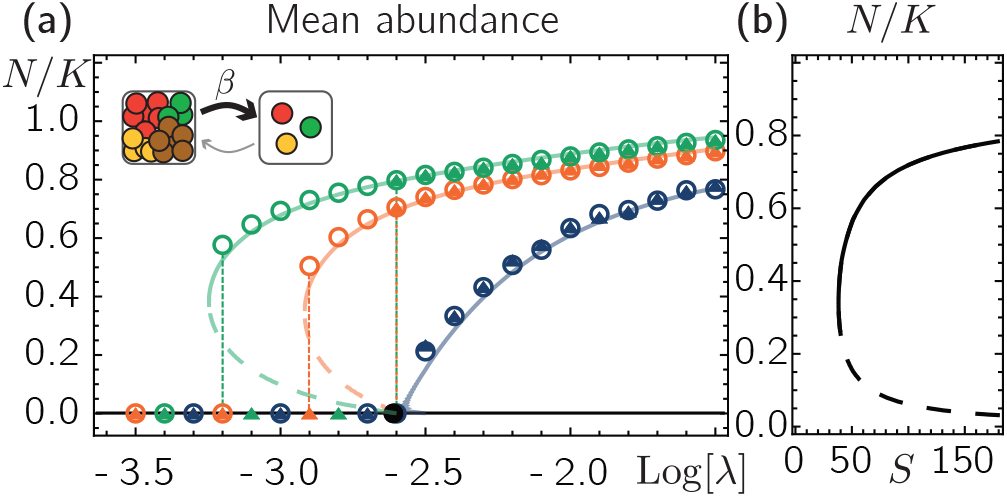
Density-dependent dispersal generates tipping point. **(a)** The mean population size in our numerical solutions for *S* =1 (blue), *S* = 40 (orange), and *S* = 100 (green), can reach different values when starting at small and large initial population sizes (triangles and circles, respectively) in agreement with our mean-field solution (solid and dashed lines denote stable and unstable manifolds, respectively). Full black circle marks *λ_c_*. The inset illustrates density-dependent dispersal, where emigration increases with the abundance of individuals from other species on a patch. **(b)** Mean-field solution of the mean abundance predicts a catastrophic shift as function of the species number, *λ* = 0.001. Remaining parameters: *r* = 0.3, *K* =10, *β* = 0.02, *P* = 500.

## Discussion

While strong (obligate) mutualism with inherent Allee effects has repeatedly been shown to yield rich behavior also in a spatially extended system, we find here that even the weakest form of mutualism among species can lead to dramatic consequences in a metacommunity that would remain hidden in a well-mixed deterministic description. Specifically, weak mutualism can have severely affect the stability of a metacommunity and lead to tipping points and hysteresis. The emergent tipping point predicts sudden shifts between coexistence and metacommunity collapse when parameters such as the dispersal rate, the number of interacting species or the strength of mutualistic interactions are even slightly varied, e.g. due to (environmental) perturbations. The emergence of bistability in our stochastic description may seem unexpected on the background of the smoothing effect of demographic noise [(30, 32) and Appendix 3], and highlights the multifaceted role of demographic noise for the dynamics of spatially extended metacommunities.

Our results pinpoint the important role of mutualistic interactions also in complex metacommunities with random interaction statistics. We show that, in general, community assembly selects for mutualistic interactions, and for dispersal rates below a critical rate *λ_c_*, mutualism is even essential to avoid metacommunity collapse. Based on these insights, our approach advocates for simplistic models that can help to understand isolated features of complex ecological systems and may offer a more intuitive interpretation.

We found qualitatively similar effects when we allowed for positive density-dependence in the dispersal rates, which suggests a more general view of mutualistic interactions.

## ACKNOWLEDGEMENTS

While completing this work, we became aware of Giulia Lorenzana, Giulio Biroli and Ada Altieri working on metapopulations with random interactions, Eq. (3), using replica methods. We thank them for stimulating discussions about our complimentary approaches. We thank current and former members of the Hallatschek lab for helpful comments and discussions. O. H. acknowledges support by a Humboldt Professorship of the Alexander von Humboldt Foundation. This research was supported by a National Science Foundation CAREER Award (1555330) and used resources of the National Energy Research Scientific Computing Center, a DOE Office of Science User Facility supported by the Office of Science of the U.S. Department of Energy under Contract No. DE-AC02-05CH11231 using NERSC award BER-ERCAP0024898. J. D. acknowledges support from the Deutsche Forschungsgemeinschaft (DFG, German Research Foundation) through grant 445916943.

## Supplementary Note 1: Numerical solution of the metacommunity dynamics

As detailed in the main text, the metacommunity is assumed to follow the dynamics Eq. (1). By rescaling the growth rate and the dispersal rate with the rate *ω*, we measure time in units *ω*^-1^ and can set *ω* = 1 in the following. Unless noted otherwise, for Fig. 2 and 3 in the main text we fixed the growth rate (*r* = 0.3), the competition strength (*α* = 0.005), the carrying capacity (*K* = 10), and solved the dynamics for global dispersal for various dispersal rates *λ* and different numbers *S* of initially coexisting species based on the following Euler forward scheme. All calculations were performed in `Python` (45) and the results were evaluated using `Mathematica` (46). For each time step Δ*t*, we first calculate the update of the population size for each species on each patch given by the growth and dispersal dynamics [first two terms in Eq. (1), respectively]. The contributions from growth and dispersal are calculated and updated separately in order to avoid unphysical scenarios, e.g. that an unoccupied patch acts as a source of dispersal. After updating the deterministic abundance of each species on each patch, demographic fluctuations [last term in Eq. (1)] are added by sampling from a Poisson distribution with the mean being the deterministic abundance. Interpreting Eq. (1) in the Itô sense (47), the Euler forward update for the demographic fluctuations are then incorporated by adding 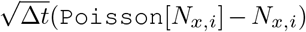 to the deterministic abundances, where `Poisson`[*N_x,i_*] is a sample from a Poisson distribution with mean *N_x,i_*. The implementation of demographic fluctuations through a Poisson process guarantees that their variance is given by *N_x,i_* (47), consistent with Eq. (1). As initial condition we either choose *N_p,i_* = 15 (large initial population sizes) or *N_p,i_* = 1 (small initial population sizes) for all patches *x* ∈ {1,...*P*} and species *i* ∈ {1,...*S*} with small random fluctuations. For the numerical solution of the more general metacommunity given by Eq. (3) in the main text we employ an analogous Euler forward scheme as above where the interaction strengths *α_i,j_* are drawn from normal distributions centered around *μ*, with standard deviation *σ/S*. For our numerical solutions, the time steps Δ*t* are adapted to values between 0.02 and 1 and the last time step is chosen to be 20000, measured in units *ω*-1. The Python code developed for this study is based on the code of our previous study on metacommunities with competitive interactions available at https://github.com/Hallatscheklab/Self-Consistent-Metapopulations, where we account for mutualistic interactions by changing the sign of the interaction parameter *α* in the code.

## Supplementary Note 2: Self-consistent mean-field approach

In the following we will discuss our analytical mean-field approach to species-rich metacommunities with global dispersal that allows us to calculate static quantities such as the critical dispersal rate *λ_c_*, the mean population size of species, and the abundance distribution. For convenience, we describe our analysis based on the dynamics for the relative abundances *f_x,i_* = *N_x,i_*/*K*. and introduce the mean fields 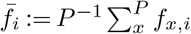 and 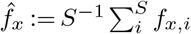 where 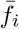 and 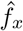 denote the averages of the relative abundance *f_x,i_* taken over patches and species, respectively. In the following we assume that the number of patches *P* and the number of coexisting species on each patch are large, and that all species are statistically identical [with identical growth rates, carrying capacities, interactions and dispersal rates, as in Eq. (1)]. Under this assumption, we estimate the sums over different patches and species in Eq. (1) through the mean field expressions 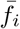 and 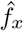 and treat these mean fields as deterministic parameters. In the following, we will first present the mean-field description for the case of direct mutualistic interaction as in Eq. (1). Later, we discuss an analogous solution for density-dependent dispersal with a dispersal term introduced in Eq. (4).

### Mean-field approach for direct mutualistic interactions

Expressing Eq. (1) through the mean fields 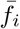 and 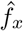, the dynamics for every species on every patch can be written as

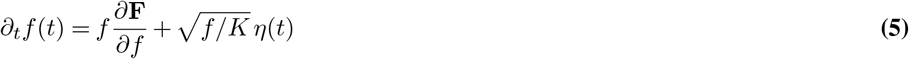

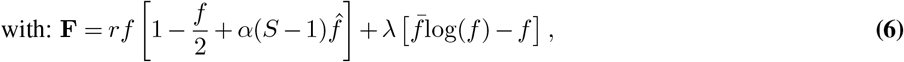

where we omitted the species and patch index since all species and patches are assumed statistically identical. The representation Eq. (5) admits an analytical equilibrium distribution in terms of a Gibbs measure (20, 48–50). To see this, it is convenient to introduce the variable 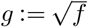. Using Itô’s lemma, we can rewrite Eq. (5) in terms of *g* as

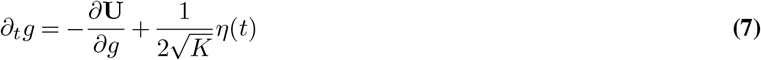

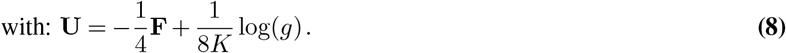

This dynamics for *g* can be reinterpreted as the overdamped dynamics of a particle in a potential **U** with diffusion constant 1/(4*K*). The equilibrium distribution 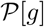 for *g* is then given through the Gibbs measure

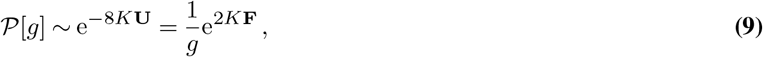

which is equivalent to a Boltzmann distribution with an ‘‘energy” given by **F**.

In terms of the relative abundance *f*, we have 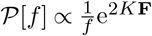, which can be expressed more conveniently as

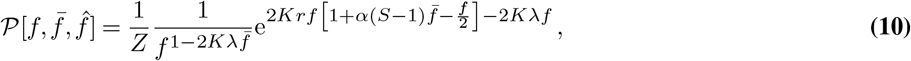

where we omitted the dependence of 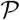 on *r, K, α*, and *λ*, and *Z* denotes the normalization constant. In terms of the abundance *N = Kf*, the distribution can be written as

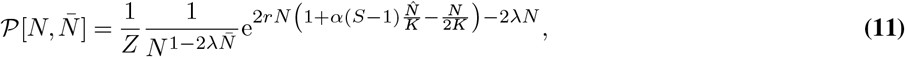

with respective normalization *Z*. While we treated the mean fields 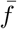 and 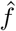 as deterministic parameters, in order for our analysis to be self-consistent they have to be equal and also equal the actual statistical mean of *f*, which can be calculated from the distribution Eq. (10). Introducing a Lagrange multiplier +*ϵf*/2*K* into the function F, we can take the derivative of log(*Z*) w.r.t to *ϵ*, take the limit *ϵ* → 0, and thereby obtain the mean abundance 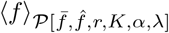. Self-consistency then requires:

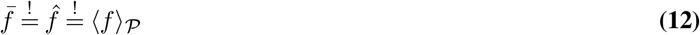

Fig. 5(a) shows the calculated mean 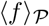 as a function of 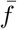 (where for specificity *r* = 0.3, *K* = 10, *α* = 0.1, and 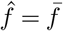 due to self-consistency). All calculations were performed using `Mathematics` (46). Varying the dispersal rate *λ* we find that for small *λ* the only solution to the self-consistency condition, Eq. (12), is given by 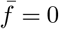. For *S* = 1, increasing *λ* above a critical value *λ_c_*, the solution 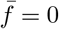 is no longer stable; however, there appears a second solution with non-zero 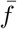, which is linearly stable and increases with *λ* [see Fig. 5(a)]. Thus, *λ_c_* marks a bifurcation from zero to non-zero mean abundances. For sufficiently large mutualistic interactions (e.g. *S* is sufficiently large for constant *α*), the self-consistency condition 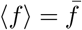 suggest an abrupt jump from zero to finite values close to the tipping point [see Fig. 5(b)].

To obtain an analytical expression for the **critical dispersal rate** we expand the calculated mean 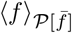 to first order in 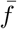. This yields the condition for the onset of finite mean abundance:

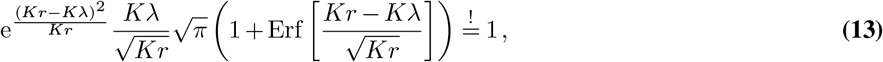

where Erf[·] denotes the Error-function (incomplete Gaussian integral). Note that the onset of finite mean abundances, Eq. (13), does not depend on the interaction strength *α* nor the number of interacting species *S*. This is consistent with the expectation that at the onset of finite mean abundances, interactions between species on a patch should be negligible. For the limiting case *K*λ ≪ *Kr*, we expand the condition Eq. (13) up to first order in *Kλ*/(*Kr*) and solved for *λ*, which yields the critical dispersal rate *λ_c_*:

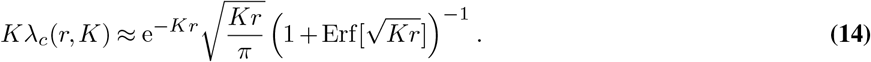

When furthermore *Kλ* ≪ 1, the growth rate must be correspondingly large so that we can set 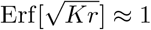. This yields 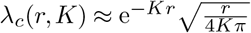. The observation of a finite dispersal threshold for global dispersal is consistent with previous studies of metapopulations with implicit spatial extension (24–26), which used master equations to model species birth, death and global dispersal between patches through a shared reservoir. In the limiting case *Kλ* > *Kr* we can expand the condition Eq. (13) to leading order of large *Kλ*/(*Kr*), and solve for *λ*. This yields the approximation:

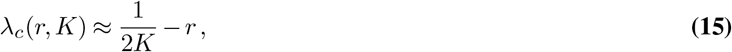

Hence, we find that for infinitesimally small finite growth rates *r*, the critical dispersal rate approaches *λ_c_*(*r,K*) = 1/(2*K*). Both limiting behaviors of the critical dispersal rate *λ_c_*, at *Kλ*/(*Kr*) ≪ 1 and *Kr*/(*Kλ*) ≪ 1, are in very good agreement with respective numerical solutions of the Langevin equation Eq. (1) [see Fig. 5,(c)].

Beyond the onset of finite mean abundances (*λ* > *λ_c_*), we can solve for the mean abundance 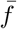 that satisfies the self-consistency condition Eq. (12) numerically [see Fig. 5(a),(b)]. Eventually, substituting this numerical solution for 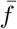 into Eq. (10) yields the *equilibrium abundance distribution* 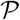 as a function of *r, K, α*, and the dispersal rate *λ*. The derived abundance distribution is governed by different contributions, depending on the choice of parameters: When the dispersal rate is small 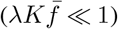 and for *f* ≪ 1 (i.e. abundances *N* ≪ *K*), the distribution of the relative abundance *f* follows the scaling

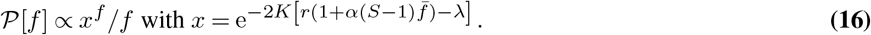

**Fig. 5.**
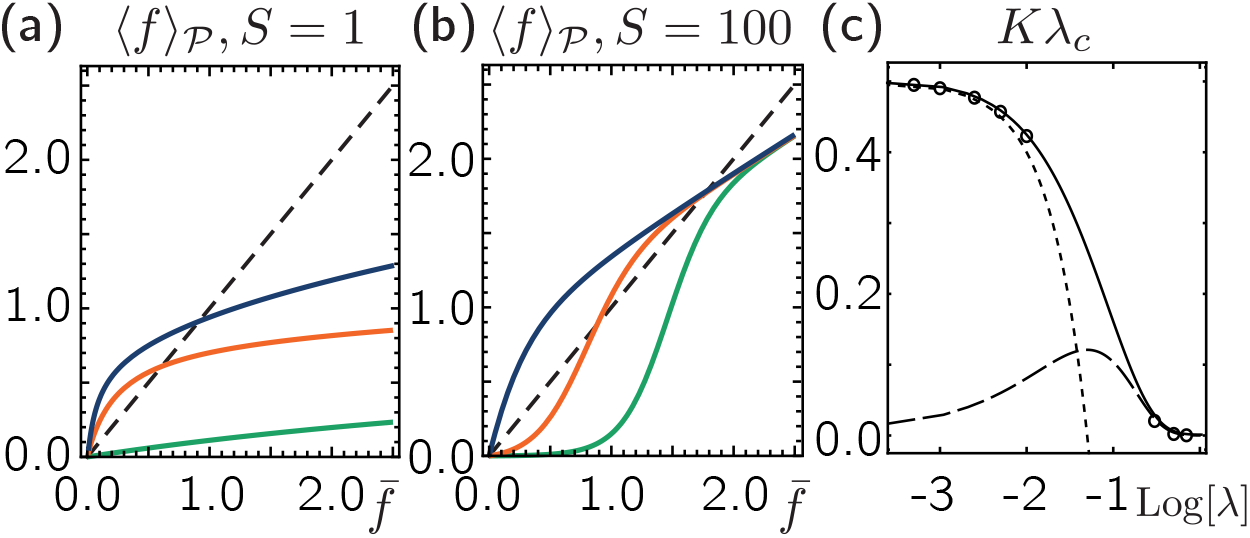
Self-consistency condition in the mean-field approximation. **(a)** Above a critical dispersal rate, *λ_c_*, the self-consistency condition 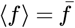 (dashed line) has a solution with non-zero mean abundance 〈*f*〉 marking an onset of finite population sizes. Shown are 〈*f*〉 for dispersal rates smaller (green, *λ* = 10^-3^), close above (orange, *λ* = 10^-2^) and farther above (blue, *λ* = 10^-1^) the critical dispersal rate *λ_c_*. **(b)** For sufficiently large mutualistic interactions (here, *S* = 100, *α* = 0.005), there is a discontinuous transition at the tipping point from a solution with zero to a solution with finite mean abundances. Shown are 〈*f*〉 for dispersal rates smaller (green, *λ* = 10^-5^), close above (orange, *λ* = 10^-3.5^) and farther above (blue, *λ* = 10^-2^) the tipping point. Remaining parameters: *r* = 0.3, *K* = 10. **(c)** The self-consistent mean-field solutions for the critical dispersal rate *λ_c_*(*r*) (solid line) are in very good agreement with our numerical solutions. The dashed and dotted lines denote the limiting behaviors for *Kλ/*(*Kr*) ≪ 1, Eq. (14), and *Kr/*(*Kλ*) ≪ 1, Eq. (15), respectively. Remaining parameters: *K* = 10, *S* = 1, and for the numerical solutions: *P* = 2000.

The form of the abundance distributions 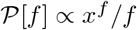 is well-known in ecology literature as Fisher log series (51), which denotes one of the most widely used abundance distributions in ecology and has been recovered in a variety of ecological systems (see (28, 29) for reviews). For larger relative abundances (*f* ~ 1), the exponential term in Eq. (10) suggests a local maximum of the abundance distribution characterized by a Gaussian distribution with mean 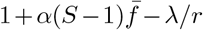 and a variance 1/(2*Kr*).

### Mutualism through density-dependent dispersal

As detailed in the main text, we investigate mutualism through density-dependent dispersal based on a dispersal rate given in Eq. (4). Using the mean fields 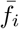 and 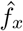, the dynamics for every species on every patch can be written as

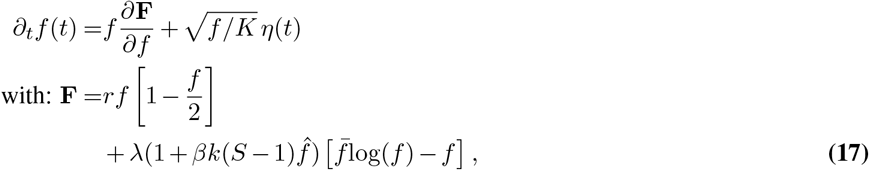

where, as in Eq. (5), we omitted the species and patch index since all species and patches are assumed statistically identical. As discussed for direct mutualistic interactions, we can use this representation to derive the equilibrium probability distribution 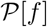 as a function of the mean fields:

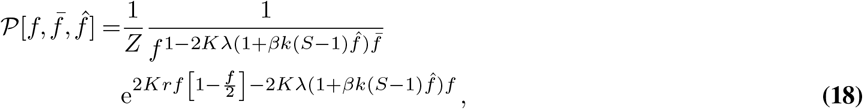

where we omitted the dependence of 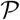 on *r,K,α*, and *λ*, and *Z* denotes the normalization constant. Imposing self-consistency, Eq. (12), we can calculate 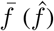 numerically and derive a closed solution for 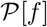. For sufficiently large *S* or *β*, the self-consistent solutions for 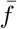 display a bifurcation depicted in Fig. 4 including a discontinuous transitions and hysteresis upon varying the baseline dispersal rate *λ*. We comment, that at the critical dispersal rate *λ_c_*, species interactions, also through dispersal, are negligible and thus *λ_c_* for mutualism through density-dependent dispersal is the same as for direct mutualistic interactions.

## Supplementary Note 3: Smoothening effect of stochastic fluctuations in metapopulations with global dispersal

One of our main results is that in the presence of mutualistic interactions between species, demographic noise can cause a sudden (discontinuous) transition between a regime where all species are extinct (inactive phase) and a regime of positive mean population size (active phase). Previous studies (30–32) have suggested that demographic fluctuations can also have the reverse effect, i.e. turn a discontinuous into a smooth transition. In their description, the population dynamics on a single patch shows bistability, with a stable fixed point at extinction and a second stable fixed point at a finite population size, separated by an unstable fixed point at intermediate population size [Allee threshold, compare Fig. 1(a)]. In Ref. (30, 32), the authors find that for short-range dispersal between patches (diffusive motion) demographic fluctuations can change the transition between an inactive and active phase from discontinuous to continuous when the dispersal rate between patches is low, or stochastic fluctuations are strong. In the following we investigate these findings for the case of global dispersal on the basis of our meanfield approximation and numerical simulations. To this end, we first describe the dynamics of a single population with size *N* that displays an Allee effect as

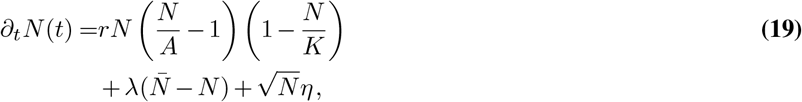

where we omitted the patch index to facilitate notation. This dynamics features bistability as illustrated in Fig. 1(a), with the Allee threshold (unstable fixed point) given by *A*. Analogous to Appendix 2, we employ a mean-field approximation, where the first term of the function **F** defined in Eq. (5), which accounts for the population dynamics, is now given by 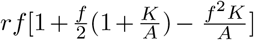. Figure 6(a) shows the obtained mean-field solutions for the mean population size 〈*N*〉 for *r* = 0.3, *K* =10 and *A* = 2 and a varying dispersal rate *λ* (lines), together with respective numerical solutions (triangles and circles denote simulations starting at small and large initial mean population sizes, respectively). As expected, for very small dispersal rates, the population goes extinct, while for larger dispersal rates there is a stable solution with positive population size. Interestingly, for intermediate dispersal rates, there is a single stable solution, while for larger dispersal rates, the system either approaches extinction or a finite population size, depending on the initial population size. Our mean field approach reveals an unstable solution which separates these two stable outcomes. This strongly suggests that, depending on the dispersal rate, the metapopulation features a strong Allee effect (high dispersal rates) or not (intermediate dispersal rates). For a better comparison with the results of (30), who plotted the mean population size as a function of the linear growth rate, we will adapt our notation in the following to their description and investigate a possible change from an abrupt to a smooth transition as they observed for short-range dispersal. Analogous to (30), we define the population dynamics

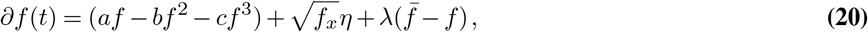

where, in contrast to (30), we included global dispersal (last term) instead of diffusive motion. *η* denotes Gaussian (white) noise with zero mean and a variance we set equal to 1 in the following for simplicity. For negative *b* and *a* and positive *c*, the deterministic dynamics of Eq. (20) suggests bistability as illustrated in Fig. 1(a), characterizing an explicit strong Allee effect. Following (30), we now fix the parameters *b* = —2 and *c* =1, and vary the parameter *a* for different dispersal rates *λ* [see Figure 6(b)]. Similar to what (30) observed for short range dispersal, when increasing *a*, we see that low dispersal rates promote a discontinuous transition from zero to finite mean population sizes, while for larger dispersal rate, the transition is smooth (i.e. continuous). Together with our results discussed above [Fig. 6(a)], this strongly suggests that also for global dispersal, a population dynamics that features bistability in terms of a strong Allee effect, can, depending on the growth and dispersal parameters, result in a smooth transition between a regime of extinction and finite mean population sizes when embedded in a metacommunity undergoing demographic noise.

## Supplementary Note 4: Metacommunities with random interactions

For random interactions, we showed that the mean interactions of survivors *I*, defined in the main text as *I* = 〈(*α_i,j_*/*K*)*N_j,x_*〉_*x,j*_, are positive in the intermediate dispersal rate regime of bistability and can be negative in for larger dispersal rates beyond the bistability regime [see Fig. 3(b)]. While in Fig. 3(b), we joined all calculated mean interactions *I* from simulations with three different sets of random interactions, Fig. 7(a) shows the distributions of *I*, 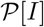, plotted separately for each set of random interactions (each in a different color).

To investigate how the communities of remaining survivors of our numerical solution and their mean interaction compare to a mean-field approximation with a species-independent interaction coefficient, we plot the mean-field solution taking into account only the number of surviving species and their average interaction coefficient *α* = 〈*α_i,j_*〉 (where the average is taken only over surviving species *i* and *j*). Interestingly, we observe that in the bistability regime [see Fig. 3(a)], the resulting mean interaction coefficient puts the community very close to the tipping point of our mean-field solution [Fig. 7(b)]. This suggests that random metacommunities self-organize to a state very close to the tipping point where they are very fragile against perturbations e.g. in the dispersal rate and species number.

**Fig. 6.**
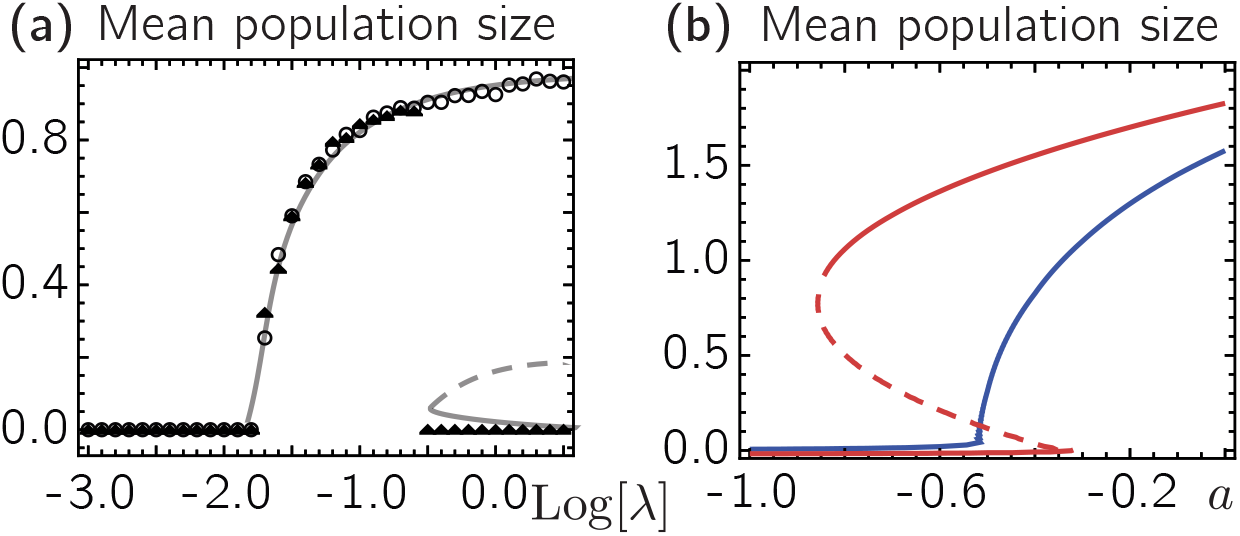
Smooth and discontinuous transitions in metapopulations with explicit strong Allee effect. **(a)** Our mean-field approach (lines) and numerical solutions (circles and triangles correspond to simulations with initially large and small mean population size, respectively) of a metapopulation with explicit strong Allee effect, Eq. (19), suggests bistability for large dispersal rates and a smooth transition from extinction to a finite population size for small dispersal rates. Remaining parameters: *K* = 10, *A* = 2, *r* = 0.3, for the numerical solutions: *P* = 1000. **(b)** Similar to what (30) observed for short range migration, we find a change from a smooth transition for small dispersal rates (blue, *λ* = 0.3) to discontinuous transition with a regime of bistability for large dispersal rates (red, *λ* = 3). The dashed line denotes unstable solution.

**Fig. 7.**
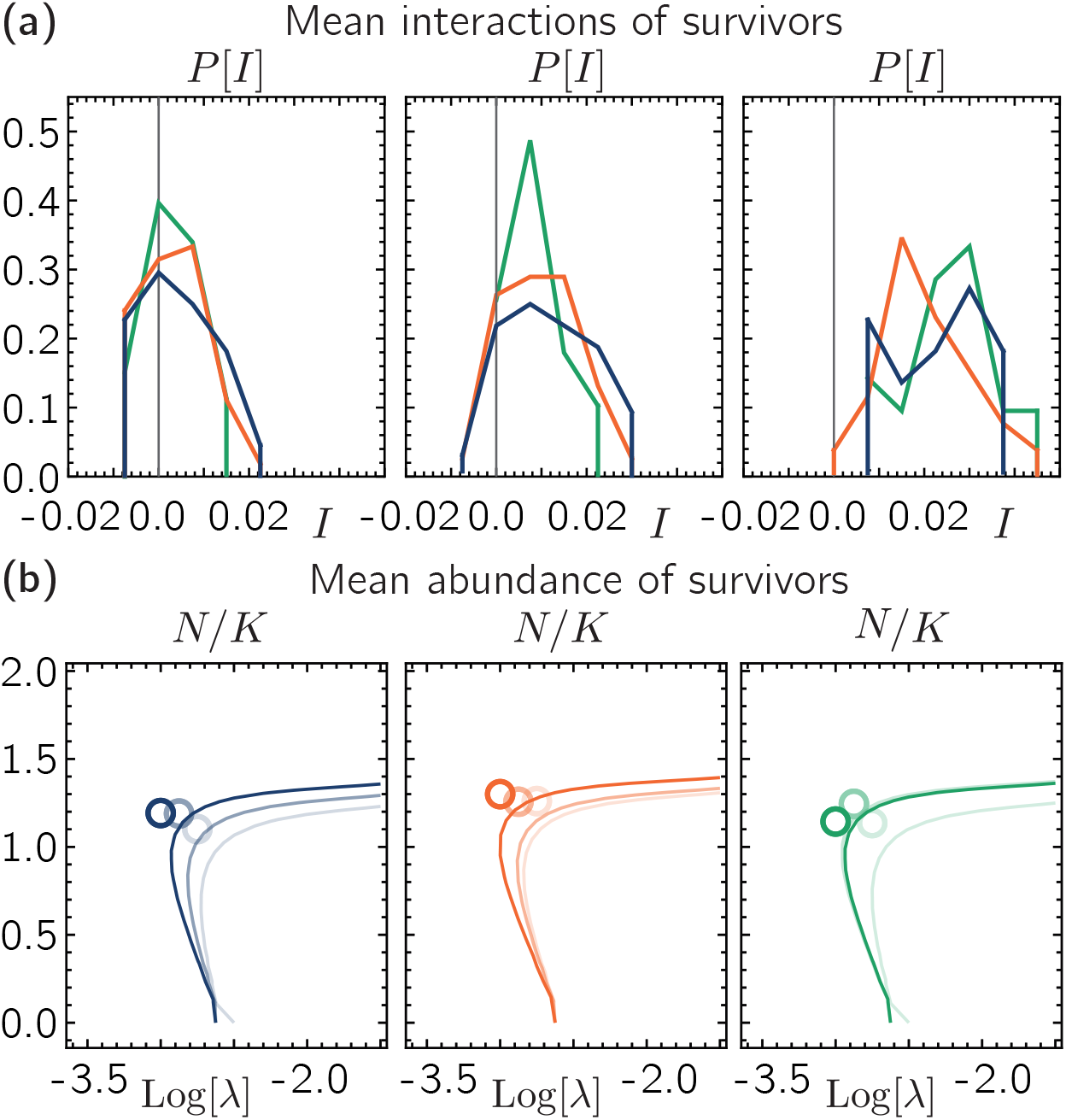
Strong metacommunity Allee effect and self-organized tipping points in random metacommunities. **(a)** Distributions 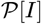 for individual independent sets of random interactions as shown jointly in Fig. 3(b) for *λ* = 10^-1.5^, *λ* = 10^-2.2^, and *λ* = 10^-2.8^ (from left to right). Different colors correspond to the three different sets of interaction coefficients *α_i,j_* used in in Fig. 3(a). **(b)** Each panel shows the measured mean abundances of surviving species for three different dispersal rates in the regime of bistability shown in Fig. 3(a). Dark to bright colors correspond to *λ* = 10^-3^, *λ* = 10^-2.875^, and *λ* = 10^-2.75^, respectively; the three panels correspond to the three sets of random interaction coefficients used in (a) and Fig. 3. Mean-field solution calculated based on the number of surviving species (ranging from ~ 25 at *λ* = 10^-3^ to ~ 55 at *λ* = 10^-2.75^) and mean of the interaction coefficients among the surviving species for the corresponding three different dispersal rates (dark to bright colors correspond to *λ* = 10^-3^, *λ* = 10^-2.875^, and *λ* = 10^-2.75^, respectively). Remaining parameters: *S* = 100, *K* = 10, *r* = 0.3, for the numerical solutions: *P* = 500.

## Footnotes

1 A weak Allee effect, in turn, refers to scenarios where at low population size, the growth rate of a species increases with its population size but stays positive.

